# Unsupervised identification of significant lineages of SARS-CoV-2 through scalable machine learning methods

**DOI:** 10.1101/2022.09.14.507985

**Authors:** Roberto Cahuantzi, Katrina A. Lythgoe, Ian Hall, Lorenzo Pellis, Thomas A. House

## Abstract

Since its emergence in late 2019, SARS-CoV-2 has diversified into a large number of lineages and globally caused multiple waves of infection. Novel lineages have the potential to spread rapidly and internationally if they have higher intrinsic transmissibility and/or can evade host immune responses, as has been seen with the Alpha, Delta, and Omicron variants of concern (VoC). They can also cause increased mortality and morbidity if they have increased virulence, as was seen for Alpha and Delta, but not Omicron. Phylogenetic methods provide the gold standard for representing the global diversity of SARS-CoV-2 and to identify newly emerging lineages. However, these methods are computationally expensive, struggle when datasets get too large, and require manual curation to designate new lineages. These challenges together with the increasing volumes of genomic data available provide a motivation to develop complementary methods that can incorporate all of the genetic data available, without down-sampling, to extract meaningful information rapidly and with minimal curation. Here, we demonstrate the utility of using algorithmic approaches based on word-statistics to represent whole sequences, bringing speed, scalability, and interpretability to the construction of genetic topologies, and while not serving as a substitute for current phylogenetic analyses the proposed methods can be used as a complementary approach to identify and confirm new emerging variants.

---

The rapid spread of SARS-CoV-2 during 2020 and over subsequent years resulted in major healthcare and societal challenges at the global level. As the COVID-19 pandemic developed, the periodic emergence of new variants that are more transmissible and/or escape host immune responses has given rise to repeated waves of infection that have each produced considerable burdens of disease despite high rates of vaccination and prior infection history [34]. There is now extensive effort in identifying worrying new variants at the very earliest stages of their emergence, which at best may enable elimination of these variants before they become established [32], but otherwise enables forward planning that may, in the future, include the timely production of tailored vaccines.

SARS-CoV-2, like other RNA viruses, has a high mutation rate and a short generation time, meaning it evolves extremely rapidly and on the same timescale as transmission. As a consequence, phylogenetic analysis has been a powerful approach to monitor the evolution of SARS-CoV-2 [28, 26, 9]. Most point mutations, or single-nucleotide polymorphisms (SNPs), that appear on phylogenetic trees are neutral or nearly neutral, meaning the mutations themselves have little or no impact on transmission and on whether lineages will grow or die out. However, some mutations do have a selective advantage because they enable more effective transmission in the population at the time of emergence. A hallmark of SARS-CoV-2 evolution has been the emergence and spread of variants of concern (VoC) and some of their major sublineages, with each having a large collection, or constellation, of lineage-defining mutations, many of which have given these viral lineages transmission advantages that have enabled them to spread extremely effectively [4].

Identifying viral lineages that are likely to be problematic in the future requires considerable effort [28] on tasks such as: the alignment of sequences to a reference before their incorporation into a phylogenic tree; the designation of new lineages (namely giving them a nomenclature based on their clade location); and the identification of lineages with potentially troublesome mutations or that are expanding quickly [9]. The production of phylogenies containing all high-quality SARS-CoV-2 genomes could help in the automation of this process, but with nearly 16 million sequences available in the GISAID database as of July 2023 (and growing), aligning a significant fraction of the sequences and generating a single phylogeny is only possible with extremely large computational resource and by making strong parsimony assumptions [35, 6]. Alignment-free methods potentially offer an option for designating new lineages that is less computationally exhausting and can cope with massive amounts of genomic data [13, 44, 43, 36, 41, 10, 16, 2].

Alignment-free techniques to characterise sequences can be classified into two types: information-theory and word-statistics based. Information-theoretic approaches rely on the notion of entropy or uncertainty between sequences, whereas word-statistic approaches, which are the focus of this project, measure the probabilistic properties of “words” (i.e. substrings) within a single sequence. An advantage of word-statistics methods, which have been used previously for interspecies genetic analyses [17, 39, 18, 1, 10], is the representation of each sequence as a vector of *n* numbers independent from other sequences. This means that distances between sequences can be calculated from their position in *n*-dimensional space ℝ^*n*^ rather than pairwise.

One of the most widely used word-statistics is the k-mer count (*kmc*, also known as k-words count), which is the count of the occurrence of words of length *k* formed by all possible combinations of an alphabet (in this case nucleotides) in a sequence; a more mathematical definition can be found in the Materials and methods section. Another featuring method that could also be considered word-statistics, known as natural vectors (*nv*), retains information on the distribution of words by creating an array of summary statistics values such as: count, mean distance to the first site of the sequence, and the variance of said distance. This method has inspired other forms to characterise genetic sequences which were also explored [40, 8, 19, 7, 27]. However, after a systematic comparison of the utility of these feature extraction approaches, we decided to focus on *kmc*, since it presented the best evaluation according to our chosen metrics (ARI and AMI) at a low processing cost. The analysis to compare results obtained using different featuring methods is shown in the Supplementary Material.

These word-statistic features extracted from data can then be processed using dimensionality reduction algorithms to help with the interpretability of relations among the sequence-data of sampled viruses by projecting such data onto lower dimensions. The projection onto reduced dimensions also allows improved unsupervised classification methods to identify clusters of sequences that merit further investigation. Classical phylogenetic analysis in fact involves a chain of feature extraction / dimensionality reduction, and then unsupervised (agglomerative) clustering. There are, however, multiple other approaches to these tasks, which can be selected to allow us to designate (give a new nomenclature to a given sequence based on a phylogenetic framework) and assign (provisionally label a sequence as a given lineage based on phylogenetic similarity) in a computationally efficient way, whilst enabling easy incorporation of new data.

Given the complex nature of the data, we found that methods developed recently to deal with features observed such as non-linearity and heterogeneous clusters performed best. In particular, we performed dimensionality reduction from the feature vector to 2 and 3 dimensions using PaCMAP [38], and identification of clusters / likely lineages using CLASSIX [5] and HDBSCAN [3]. These methods lack the representation of biological reality present in a phylogeny, but scale in a computationally efficient manner with data volume, so may prove to be important complementary tools to track the evolution of SARS-CoV-2 as well as other rapidly evolving pathogens.

## Results

As detailed in the Materials and Methods section we worked with 5.7 million complete high-coverage genetic sequences from GISAID [31]. To demonstrate the potential of using alignment-free features to represent the genetic structure of SARS-CoV-2 populations, we first compared a projection of *3mc* (*kmc* with *k* = 3) from unaligned sequences with a maximum parsimony phylogeny generated using IQTree [24] using an alignment of the same sequences shown in Figure 1. Because of the computational costs of generating phylogenetic trees, we used a stratified subsample of 5 000 high coverage sequences. The multi-dimensional *3mc* features were projected onto a 2-dimensional space using the non-linear dimension reduction tool PaCMAP [38], with each point representing the location of the sequence in the estimated genetic landscape. The points are coloured based on the Scorpio labelling by the Pangolin tool (see Materials and methods) [26]. The similarity of the clusters of the significant VoC, such as Alpha (B.1.1.7), Delta (B.1.617.2), Omicron (BA.1), and Gamma (P.1), from both the 2d PaCMAP projection and the maximum parsimony tree, demonstrates the potential of the combination of word-statistics features and dimensionality reduction tools to produce interpretable structures for the distribution of variants in a genetic space.

**Figure 1:**
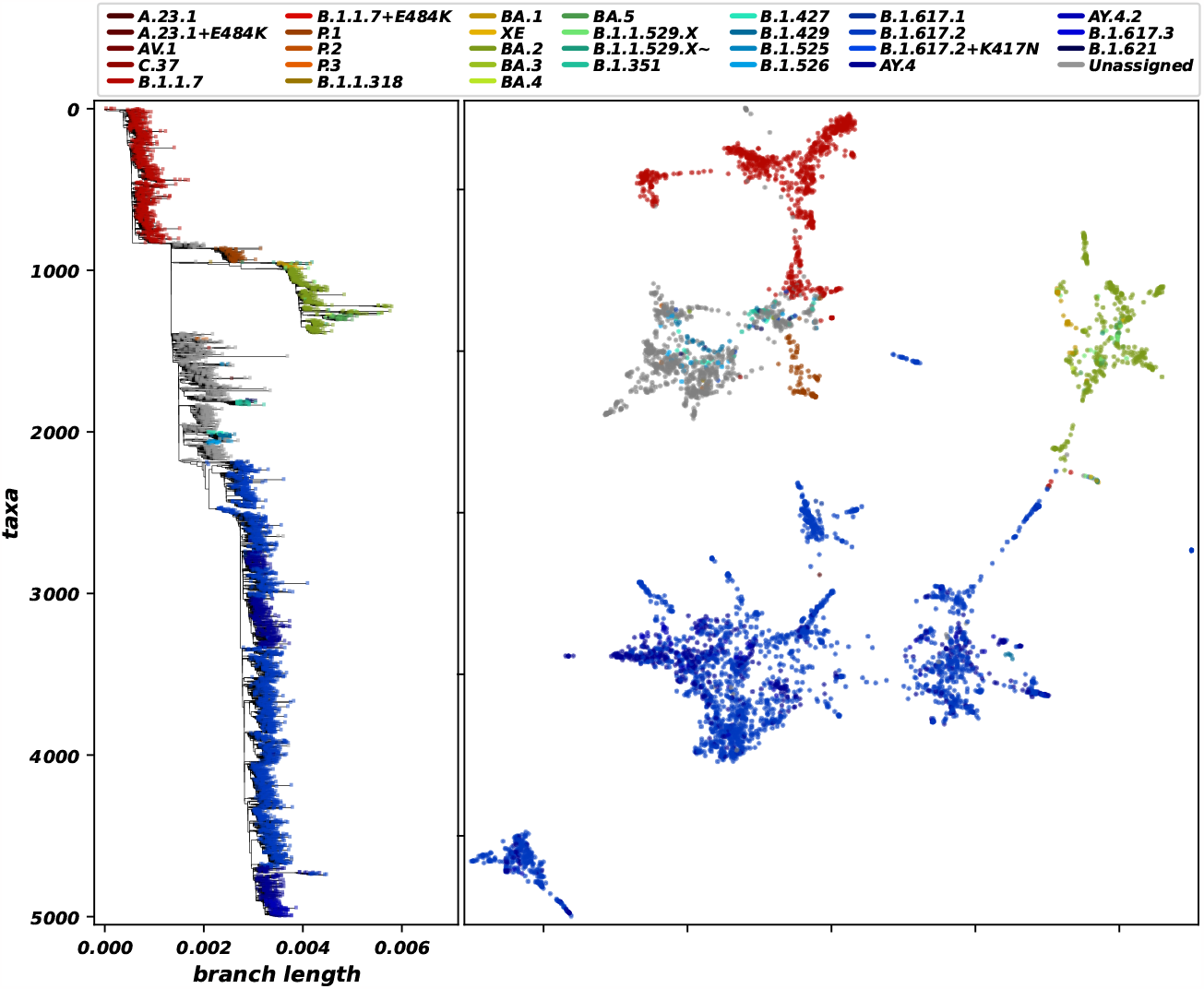
Comparison of a maximum parsimony phylogenetic tree generated using IQTree (left) and PaCMAP projection of 3-mer counts (*3mc*, right). This plot was generated using a subsampled alignment of only 5 000 high coverage genomes (*<* 1% of undefined bases with a longitude of ∼ 29*K* nucleotides) from those available on GISAID [31], up to 19 January 2023. The similarities between emerging clusters in both analyses show the potential of the combination of word-statistics features and dimension reduction tools to gain insight of the distribution of variants in a genetic space.

To measure the clusters formed by a 3-dimensional PaCMAP projection of the 5.7 million sample data quantitatively, we implemented two clustering algorithms, HDBSCAN [21] and CLASSIX [5] using a set of parameters we denote by *GISAID1* (see Materials and Methods for further details). The Scorpio labelling, produced by Pangolin, was used as a “ground truth” to calculate the adjusted Rand index (ARI) [29] and adjusted mutual information (AMI) [23] of the clustering matrices. The newer CLASSIX outperformed the more established HDBSCAN, with ARI and AMI values of 0.4741 and 0.6050 for CLASSIX, versus 0.4707 and 0.5935 for HDBSCAN. In both metrics a value of 1 would mean the emerging clusters are completely equivalent to the grouping of the “ground truth”. The results of this analysis of the *3mc* feature for the whole GISAID dataset can be seen in Figure 2.

**Figure 2:**
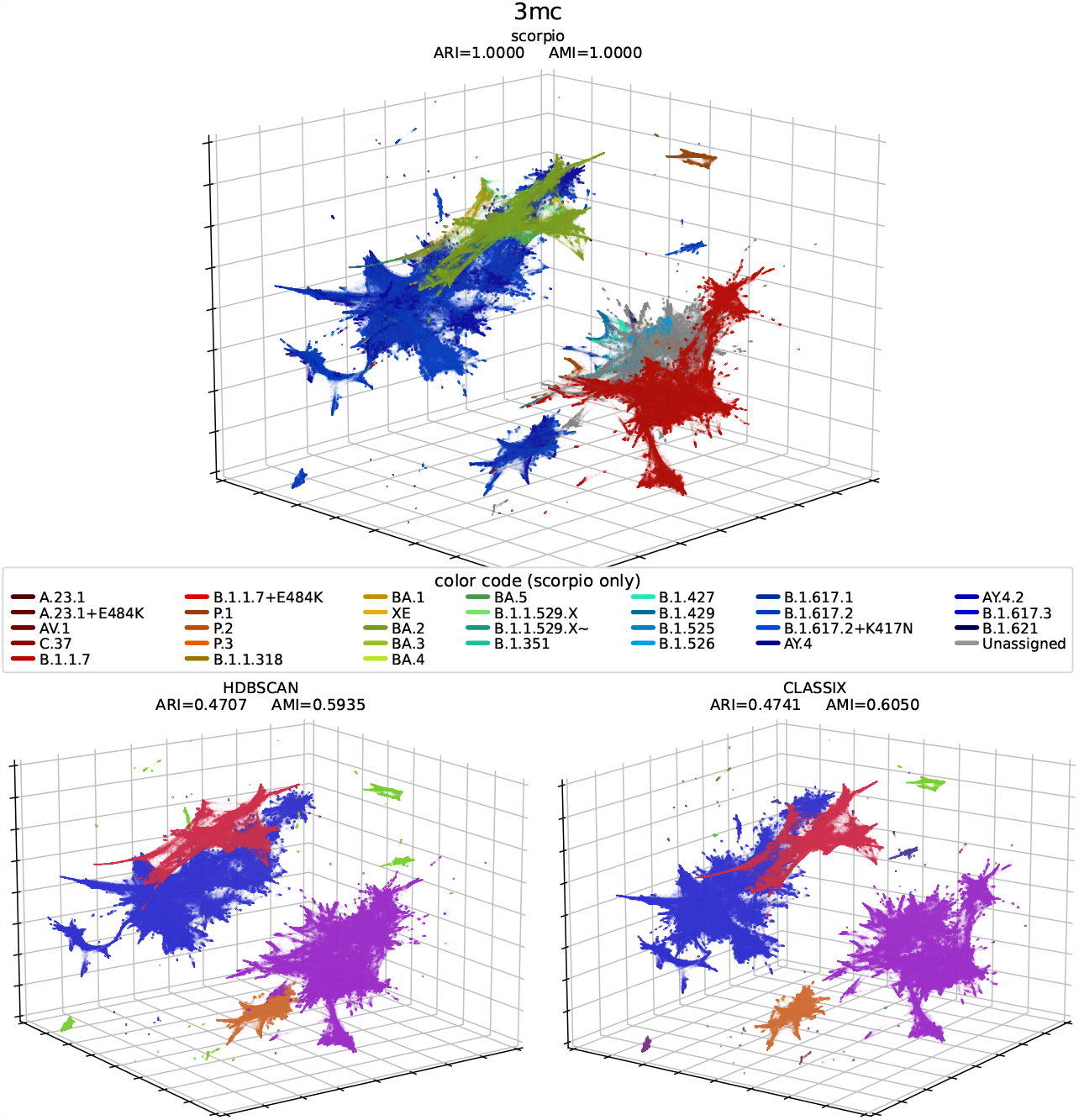
3d PaCMAP projection and clustering analysis of genetic topology of roughly 5.7 million sequences. Here we can see the extracted feature *3mc* coloured by Scorpio labelling (top, same color code as Figure 1), clustered by HDBSCAN (bottom-left), and clustered by the novel CLASSIX algorithm (bottom-right). The projection and clustering was performed using the parametric set-up *GISAID1*, detailed in Materials and Methods. In the bottom figures the color code is arbitrary, showing the unsupervised detection of clusters the algorithms perform. The formation of well defined clusters show the potential of these tools to help us gain insight on phylogenetic relations of vast amounts of sequences in a relatively computationally inexpensive fashion. Clicking on the hyperlinks in this caption leads to interactive 3d plot.

We were next interested in how the PaCMAP projection of *3mc* performed when restricted to the major structural protein regions, shown in Figure 3. To accurately identify the different gene regions we used the aligned sequences. This projection was performed using only the PaCMAP parameters of *GISAID1*, since the aim of this analysis is to compare regions of the genome rather than to identify clusters. The plots within Figure 3 show that while the S and N regions reduce into many clusters associated with different Scorpio labels, the E and M regions are much more homogeneous. The relative homogeneity observed for E and M may be due in part to their relatively short lengths (227 and 668 nucleotides respectively) compared to S and N (1 259 and 3 821 nucleotides respectively), and also possibly due to being subject to stronger adaptive evolution, in particular the existence of antibodies that bind to S and N [11].

**Figure 3:**
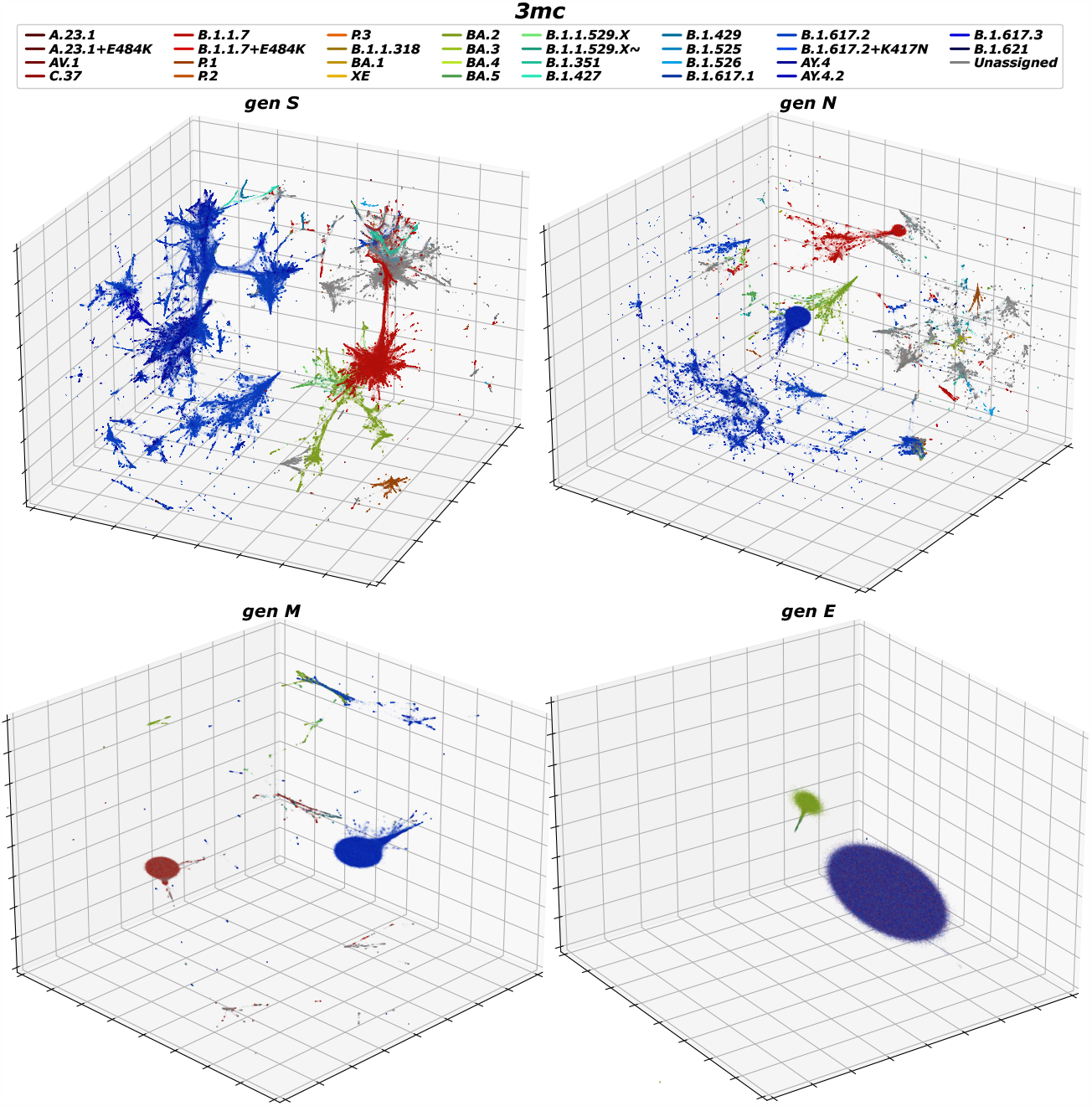
3d PaCMAP projection of specific structural protein regions for about 5.7 million sequences, generated using *3mc* and coloured by Scorpio labelling. The formation of clusters allows to see the contribution of the different proteins to the differentiation within the viral population. While proteins S and N seem scattered, arguably due to selection pressures, M and E seem to have found a stable configurations. Clicking on the hyperlinks in this caption leads to interactive 3d plots. It is worth to mention that all variants are contained within the two clusters emerging for the projections of genes M and E, however due to the high volume and mixing among them it is difficult to distinguish each of the lineages separately giving the impression of being coloured outside of the initial colour coding.

We tested the use of this pipeline to detect the emergence of new variants under the hypothesis that they will form new clusters as they appear. For this we took a subsample of the sequences collected in England (n=982 496), from which subsets were taken forming a temporal cumulative progression with 2-week steps. The results can be seen in Figure 4. There were some adjustments made to the parameters of the HDBSCAN clustering algorithm to increase the sensibility of finding small clusters, while CLASSIX parameters were left as in *GISAID1*. This new parametric set-up was called *GISAID2*. The results can be seen in Figure 4: here it is possible to see that HDBSCAN follows Scorpio in number of clusters before detecting very many more, while CLASSIX stays close, although the number grows more gradually in early 2021 than Scorpio. Further exploration on the PaCMAP parameters allowed to increase the Pearson correlation coefficient (*r*^2^) between the CLASSIX cluster detection and the Scorpio labelling, from 0.7851 to 0.9029. However this change of parameters reduced the AMI value by a mean of 0.0484, namely from a mean of 0.3228 to one of 0.2743. This third parametric set-up was named *England1*.

**Figure 4:**
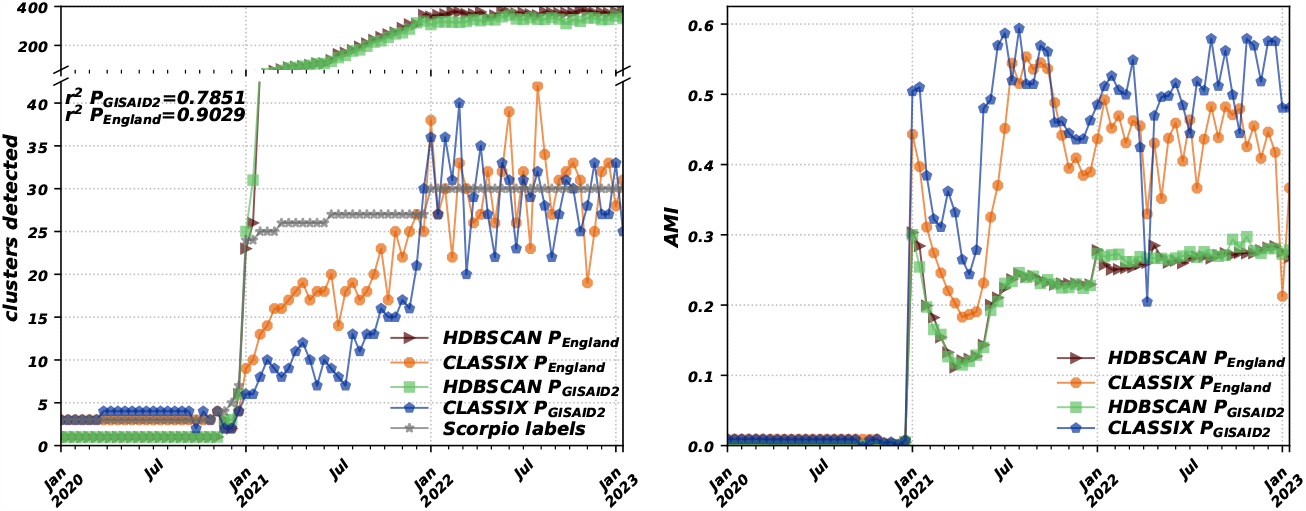
Results of an analysis applied only to the subset of GISAID sequences reported from England. The dimensionality reduction and clustering algorithms were applied to a cumulative temporal progression with resolution of two-weeks time with set-up parameters *GISAID2* and *England1* (left). The contrast between these settings showed a trade-off, such that while the *r*^2^ to Scorpio is increased from 0.7851 to 0.9029, the adjusted mutual information index is reduced from a mean of 0.3228 to one of 0.2743, which implies a reduction of the cluster quality (right). These results point to the potential of PaCMAP dimensionality reduction and clustering detection in alerting about the possible emergence of new variants by detecting the appearance of new clusters, should a regularisation and noise reduction to the cluster detection is eventually achieved.

In terms of computational demands, a rough estimate is that the whole process from feature extraction to cluster detection could be made in roughly 30 hours on a modern laptop. The *3mc* feature can process a high coverage sequence in roughly 0.12 seconds, and exploiting parallelisation the whole 5.7m-sequence dataset can be processed in about 14 hours. The PaCMAP dimensionality reduction then takes approximately 17 hours, while the clustering can be performed in less than 5 minutes. More details of the times can be seen in the Supplementary Materials.

## Discussion

The unprecedented amount of genetic data generated during the pandemic demands the development of methods to analyse it thoroughly, with fluidity and efficiency. Without showing a benefit to curating this data in the future there is a risk that it will be removed or deleted. The application of algorithms that characterise full sequences and project them onto low dimensional space has the potential to reveal genetic relationships among a huge number of sequences simultaneously. The supervision of these genetic spaces with automated clustering algorithms could provide alternative methods to discover niches of specific viral variants. Between the two clustering algorithms used, CLASSIX produced results more consistent with existing approaches (although these are not automatically completely optimal) when compared to HDBSCAN, and with a minor number of parameters it was easier to optimise. CLASSIX was even ten fold faster and in general yielded higher AMI and ARI, making it a better tool for the scope of this project. Nonetheless, more research on the parametric space exploration of HDBSCAN is needed, together with the investigation of different clustering algorithms.

The different parameters for the PaCMAP projection and of the clustering algorithms of the temporal cumulative subsets showed a trade-off, where the *r*^2^ to Scorpio increased from 0.7851 to 0.9029 but the adjusted mutual information index decreased by a mean of 0.0484, from 0.3227 to 0.2743.

This analysis serves as a proof of concept, as these results are encouraging of their potential application as an alert tool for the early discovery of an emerging VoC, or as supporting method to its discovery. Whilst phylogenetics remains the gold-standard for understanding the genetic topology of viral populations, the proposed methods have the advantage of being able to manage several orders of magnitude more sequences than the current methods at a given computational cost.

On the basis of results reported above, we expect that the whole processing of 5.7 million high-coverage sequences can be done in 1-2 days on a standard modern laptop, which fulfills the main goal of finding a feasible method to gaining insights by processing huge amounts of data. To keep the complexity low, the value of *k* for the *k* -dependent algorithms was set to 3, although we are aware that future research on this topic could deepen on the optimal values of *k* to understand the trade-offs of a proper extraction of information from the sequences and computing costs. Additionally, there could be an evaluation on which features might be most significant for a suitable construction of genetic spaces. In alternative, the application of other alignment-free methods such as “protein-*nv*” [42], graphical representation [33], and Fourier power spectrum [12], among others, could add value to these analyses. Furthermore, previous methods for phylogenetic reconstruction have used non-Euclidean distances like: Wasserstein, Kullback-Leibler, Yau-Hausdorff [33], or Structural Similarity Index Measure. Thus, applying them to dimension reduction algorithms might generate better representations of the genetic landscape. The presented results show values of ARI and AMI that seem encouraging considering the non-discrete nature of the labelling. Additionally, it has been argued [30] that these metrics are optimal for different types of clusters: in particular, ARI is advised for when the clusters present roughly equal sizes, while AMI is more appropriate for when the cluster sizes are highly unbalanced. Based on this consideration, we could argue the AMI index to be more meaningful for the presented dataset. Further analysis, like combination of features, and parametric exploration, could improve these metrics and reduce noise.

The methods presented in this paper provide the means to accelerate the identification of new pathogen variants of concern by applying alignment-free methods based on word-statistics. The extremely rapid pace of developments in algorithms and their implementation for dimensionality reduction and unsupervised clustering is expected to continue, and as such produce further valuable complementary analyses to the current phylogenetic gold-standard methods. Further exploration of hyperparameters, mixing of different methods of characterisation, and data preprocessing, could help to enhance their accuracy and reliability.

## Materials and methods

The process described in this work was run using a laptop with a processor of 11th Gen Intel(R) Core(TM) i7-11800H and 2.30GHz, with a memory RAM of 16 GB. The full GISAID database [31] was downloaded on 19 January 19 2023 when it contained 14 617 387 SARS-CoV-2 sequences. These sequences were then filtered to include only complete (*>* 29 000 nucleotides), high-coverage (*<* 1% loci identified as N) and unique sequences resulting in a total of 5 726 839. Each sequence was aligned to the reference sequence hCov-19/Wuhan/WIV04/2019 [25] using the tool MAFFT v7.453 [14] on an Ubuntu Windows subsystem for Linux v20.04.1 LTS and Biopython v1.78 scripts. Once aligned, Pangolin lineages were obtained by running PangoLEARN v1.18 [26] and taking the “Scorpio call”. To compare phylogeny with PaCMAP, as in Figure 1, a subsample of *n* = 5 000 from the high-coverage sequences dataset sequences, stratified by Scorpio lineage, was aligned and processed to generate maximum parsimony phylogenetic trees using the tools MAFFT and IQTree v2.2.2.6 [24]. This size of subsample was determined by the computational cost of phylogeny. The same subsample was then projected onto two dimensions using PaCMAP to visualise the contrast between the two techniques.

In order to extract features, we wrote bespoke code in Python v3.10.0 to extract the required word statistics and derived quantities. The only feature we consider in the main analysis – though others are explored in the supplementary material – is the k-mer count (referred as *3mc*, given *k* = 3), which is produced by sliding a window of length *k* through the whole sequence shifting one position at the time obtaining *n*− *k* + 1 overlapping strings, where *n* is the length of the sequence. From there, the occurrences of words of length *k* formed by all combinations of nucleotides are counted, resulting in a dictionary with the frequencies of each k-mer as values. This means that for a sequence AAAAACGT, this algorithm would extract a dictionary of 3-mer: {AAA : 3, AAC : 1, ACG : 1, CGT : 1}. For simplicity and to keep within the scope of scalability of this project, the value *k* was set to 3 for all *k* -dependant algorithms.

The aforementioned methods of word-statistics to characterise the sequences were then projected onto low dimensions using graph-based dimensional reduction algorithms, as a first step towards unsupervised classification. Non-linear methods such as t-SNE [20] and UMAP [22] were explored, as well as linear methods such as principal components analysis, and through this we found that PaCMAP v0.5.2.1 [38] produced the most robust and interpretable outputs.

Tuning of algorithmic hyperparameters was carried out on a 1% subset, stratified by Scorpio labelling, of the data before running on the full dataset with the selected parameters. For PaCMAP, these are the default values for projection, after which the low dimensional space was minmax normalised, namely the lowest value was normalised to 0 and the highest to 1 for all three dimensions of the projection. Then, unsupervised clustering was performed using HDBSCAN v0.8.27 [3, 21] and the novel CLASSIX v0.6.5 [5]. The parameters of these clustering algorithms were set to their defaults, with the following exceptions. For HDBSCAN, we set `min_cluster_size=200000`, since otherwise this defaults to 5, a value that generates an excessive number of small clusters in a dataset of this size. For CLASSIX, we set `radius=0.2` (compared to a default of 0.5) and `minPts=500` (compared to a default of 0).

The quality of clusters detection was then evaluated through the metrics of adjusted Rand index (ARI) and adjusted mutual information index (AMI) [37]. This exercise was repeated on a 3 *×* 3 grid for the PaCMAP parameters with combinations of the hyperparameters `MN` in {0.25, 0.5, 1.0} and `FP` in {1.0, 2.0, 4.0}, while other parameters were kept at their default values. The optimal values for the PaCMAP projection of *3mc* were found to be 1.0 and 1.0 for `MN` and `FP` respectively. We call this parametric set-up for the clustering and PaCMAP algorithms *GISAID1*. The optimal values however differ for each feature extraction method, as detailed in the supplementary material.

Additionally, to test the detection of emerging clusters through time, the PaCMAP dimensionality reduction and clustering were applied onto a series of subsets of cumulative subsampling through time. This analysis only accounts for the subset of GISAID sequences reported from England. The time resolution step was defined as 2 weeks. The PaCMAP dimensionality reduction was performed with the same parametric set-up *GISAID1*, with the exception of a change on the sensibility of the size of the cluster detection of the HDBSCAN clustering algorithm: namely, we set `min_cluster_size=500` for consistency with CLASSIX. This set-up was called *GISAID2* and resulted in an encouraging, although noisy, detection of clusters for CLASSIX matching roughly the Scorpio labels, but a tendency of HDBSCAN to overshoot its detected clusters. Further parametric exploration was made to try to match more closely the Scorpio labelling. The new parametric set-up changed the PaCMAP `MN` and `FP` parameters to 0.25 and 2.0 respectively, while keeping the parameters of the clustering algorithms the same as in *GISAID2*, and was called *England1*.

Interactive 3D plots of the PaCMAP projections, the scripts and a subsample of the dataset are available at the repository: github.com/robcah/dimredcovid19

## Acknowledgements

This project was possible thanks to the support of the Joint Universities Pandemic and Epidemiological Research Consortium (JUNIPER – https://maths.org/juniper/), the Engineering and Physical Sciences Research Council, the Wellcome Trust and the Li Ka Shing Foundation. LP gratefully acknowledges the Wellcome Trust and Royal Society (grant number 202562/Z/16/Z). TAH is supported by the Royal Society (grant number INF\R2\180067). IH is supported by the National Institute for Health Research Health Protection Research Unit (NIHR HPRU) in Emergency Preparedness and Response and the National Institute for Health Research Policy Research Programme in Operational Research (OPERA). The views expressed are those of the author(s) and not necessarily those of the NHS, the NIHR, the Department of Health or Public Health England. LP, TH and IH are also supported by the JUNIPER modelling consortium (grant number MR/V038613/1) and by the Alan Turing Institute for Data Science and Artificial Intelligence under the EPSRC grants EP/N510129/1 and EP/V027468/1. LP, TH, IH and RC also acknowledge funding from the UKRI Impact Acceleration Account (IAA 386). KL gratefully acknowledges the Wellcome Trust and Royal Society (grant number (107652/Z/15/Z)) and the Li Ka Shing Foundation.

## Supplementary Material

### Word statistics featuring methods

The viral evolution of SARS-CoV-2 in combination to the modern sequencing technologies presented a challenge for the quick analysis of the massive volume of genetic data. As mention in the main article, this project hypothesised that given that word-statistic methods were proven methods to create phylogenetical trees, the emergence of analogous structures could be appreciated through dimension reduction, thus producing an interpretable embedding showing a distribution of variants in a projected genetic space [2].

From all the summary-statistics features explained in the main paper, only *kmc* [15] results are shown in the main analysis. However, we also implemented and tested more complex methods based on natural vectors (*nv*), which we detail and present below.

The natural vectors (*nv*) approach was proposed by [7], who used standardised central moments to characterise the distribution of single nucleotides in each sequence. The natural vector of a genetic sequence *S* = (*s*_1_, *s*_2_, …*s*_*n*_) of length *m*, where *s*_*i*_ ∈ {A, C, G, T*}, i* ∈ {1, …, *n*}, is constituted by the following elements for each *ω* ∈ {A, C, G, T}:

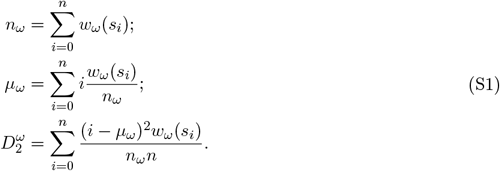

Here the count of nucleotide *ω, n*_*ω*_, the mean distance of said nucleotide to the origin of the sequence, *μ*_*ω*_, and the variance of said distance, 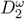, are defined in terms of the indicator functions

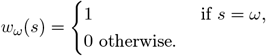

Since we have a copy of the variables in Equation [S1] for each of the four amino acids, this leads to a 12-dimensional natural vector representation of the sequence *S* that can be written explicitly as

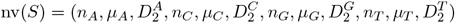

Another characterisation of sequences comes from Accumulated Natural Vectors (*anv*) [8] which forms its arrays by *n*_*ω*_ as defined in Equation [S1], together with average distance to the end of the sequence, *ζ*_*ω*_ and the covariance of the accumulation of nucleotides through *S*. These new elements can be defined as follows:

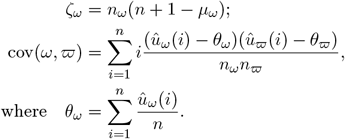

Here, *û*_*ω*_ is the accumulation indication function, which is an array of the accumulation of nucleotide *ω* in position *i*, so for *S* = AAGCGT, *û*_*A*_ = (1, 2, 2, 2, 2, 2), for *û*_*G*_ = (0, 0, 1, 1, 2, 2) and so on for all nucleotides, or formally in terms of the indicator function

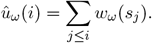

The covariance is calculated for all pairs of elements of A, C, G, T, which results in the 18-dimension array

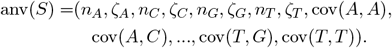

A different method called weak-amino-purine vector (or *wmrv*) [19] characterises the sequence *S* by producing three mappings of the sequence codifying it into three pairs of complementary bases. The first coding is onto purine (R) or pyrimidine (Y),

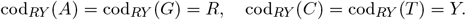

The second coding is onto amino (M) or keto (K),

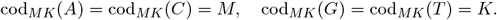

The third coding is onto strong H-bond (S) or weak H-bond (W),

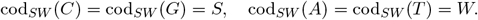

And so for *S* = AAGCGT, cod_*RY*_ (*S*) = RRRYRY, cod_*MK*_(*S*) = MMKMKK and cod_*SW*_ (*S*) = WWSSSW. The resulting array has the following 18-dimensions:

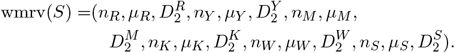

Another idea from [40] attempted to combine the use of k-mers and *nv* by characterising k-mer distributions within *S*. For this a window of length *k* is slid throughout the whole sequence, and the count, mean distance, and variance of the k-mers are characterised with the summary statistics in Equation [S1] for *nv* applied to each k-mer instead of each nucleotide i.e. with *ω* ∈{AAA, AAC, …, TTG, TTT}. This is called k-mer natural vectors, or *kmnv*, for *k* = 3.

Finally, the extended natural vector (*env*) [27] combines the frequency chaos game representation (FCGR) characterisation with the *nv* approach. In this method the objects to characterise are the discrete values *ω* ∈{0, …255} which represent the intensity of a pixel in the 2-dimensional FCGR, expressed as the square matrix *Q* with values *q*(*i, j*) ∈*ω*, where *i* and *j* are dynamic values referring to the rows and columns within matrix *Q*. The equations from *nv* are adapted to 2-dimensions as below:

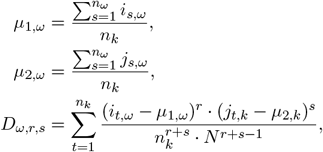

for pairs (*r, s*) ∈{(1, 1), (0, 2), (2, 0)}. Therefore the resulting 1536-dimensional array is formed by the following values:

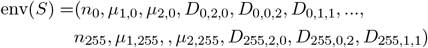

Here *k* ∈*K, r* is an arbitrary non-negative integer, and *s* = 0, 1, …*n*_*k*_, these values are to calculate the normalised central moments of the distance of the *ω* values. *D*_*k*,1,0_ and *D*_*k*,0,1_ result in zero, a property of the location of means *μ*_1,*k*_ and *μ*_2,*k*_. Thus central moments of combined degree 0 and 1 can be omitted, as well as higher central moments since converge to zero.

As seen in the following features there is mainly tendency for these characterisations to produce reasonable clusters for similar sequences. The overall projections of these natural-vectors features can be seen in Figures S4, S1, S2, S3, S5. While of interest, these do not provide qualitatively different insights from the more interpretable *kmc* feature used in the main paper.

### Dimensionality reduction

The exploration of linear algorithms for dimensionality reduction, such as PCA, ICA or SVD did not yield results with sufficient interpretability. Therefore, other non-linear algorithms were used, such as t-SNE [20] and UMAP [22]. These two had a better result on the representation of the data, though the replicability was challenging since they seemed to be very sensible to the hyperparameter settings. The novel PaCMAP [38] presented robust replicability and was therefore used to perform the analysis of this project.

This algorithm goes through three main steps: graph construction; initialisation of solution; and a three-staged iterative optimisation. The graph construction is based on distinguishing three types of edges: neighbour pairs (NB), i.e. nearest neighbors from a given observation, mid-near pairs (MN) and farther pairs (FP). Principal Component Analysis (PCA) is used to initialise the solution in the lower dimension space. From this it can be understood that the graph gives forces of attraction and repulsion to each point depending on the relationships of the high dimensional graph and the loss function *L*_PaCMAP_. The solution uses Adam Optimiser to minimise *L*_PaCMAP_ throughout iterations (*t*_1_, *t*_2_, …*t*_*q*_, where *q* = 450 as default), defined as:

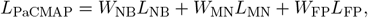

where *W*_*NB*_, *W*_*MN*_ and *W*_*F P*_ are weights varying depending the iterative stages of the algorithm. Such stages are separated by the thresholds *τ*_1_ = 1, *τ*_2_ = 101 and *τ*_3_ = 201, and the weights vary as follows:

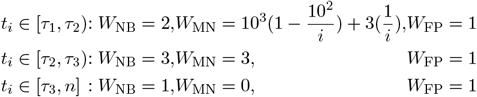

The first phase begins with heavy weights for MN pairs, which gradually decrease to preserve both global and local structures. The second phase aims to preserve the local structure by assigning a lower value to *W*_MN_. These two first stages try to prevent the lack of forces on far points and placing neighbours too far apart in early iterations, which would make it harder for these neighbours to get close in further iterations. The third stage focuses on improving local structure by setting *W*_MN_ to zero and reducing the other two weights, to emphasise the repulsive forces to separate clusters. The remaining terms are defined as:

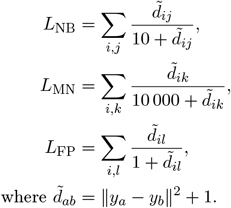

In the above *i* is the reference datapoint while *j, k* and *l* are all the points in the sets of nearest neighbour, mid-near and far datapoints, respectively. Furthermore *y* is the array of coordinates of data-points in the original *R*^*n*^ graph, with the indices ranging in their respective sets. The distance 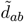 is calculated in base of the scaled squared Euclidean distance (‖ ‖ ^2^). A summary of the processing times can be seen in Figure S6.

### Clustering

HDBSCAN [3] is an extension of DBSCAN, and finds clusters by assuming an existing density function from which the datapoints are drawn, thus producing a cluster-tree considering each peak in said density function as a branch and then by “runt pruning” – focusing on branches with higher density – which is defined as groups of “core points”, namely datapoints having at least a predefined number of datapoints within a neighborhood of a predefined radius and constraining them to not overlap. This algorithm proved to be relatively fast and yielded far better results than standard methods such as k-means.

Additionally, we also tried the more recently proposed method CLASSIX [5], which has been optimised for low computational complexity and it is based on sorting the datapoints through distances on a series of principal components, starting with the first point and assigning all the points within a predefined radius to the same group, then making the same with the next unassigned datapoint. When all datapoints have been assigned to a group, they are merged into clusters if their initial datapoint of the group is found within a radius 1.5 times the aforementioned predefined distance.

### Summary

A summary of the results and processing times can be found in Table S1. For further completeness, the scripts and a subsample of the dataset are available for replication at the repository: github.com/robcah/dimredcovid19

**Figure S1:**
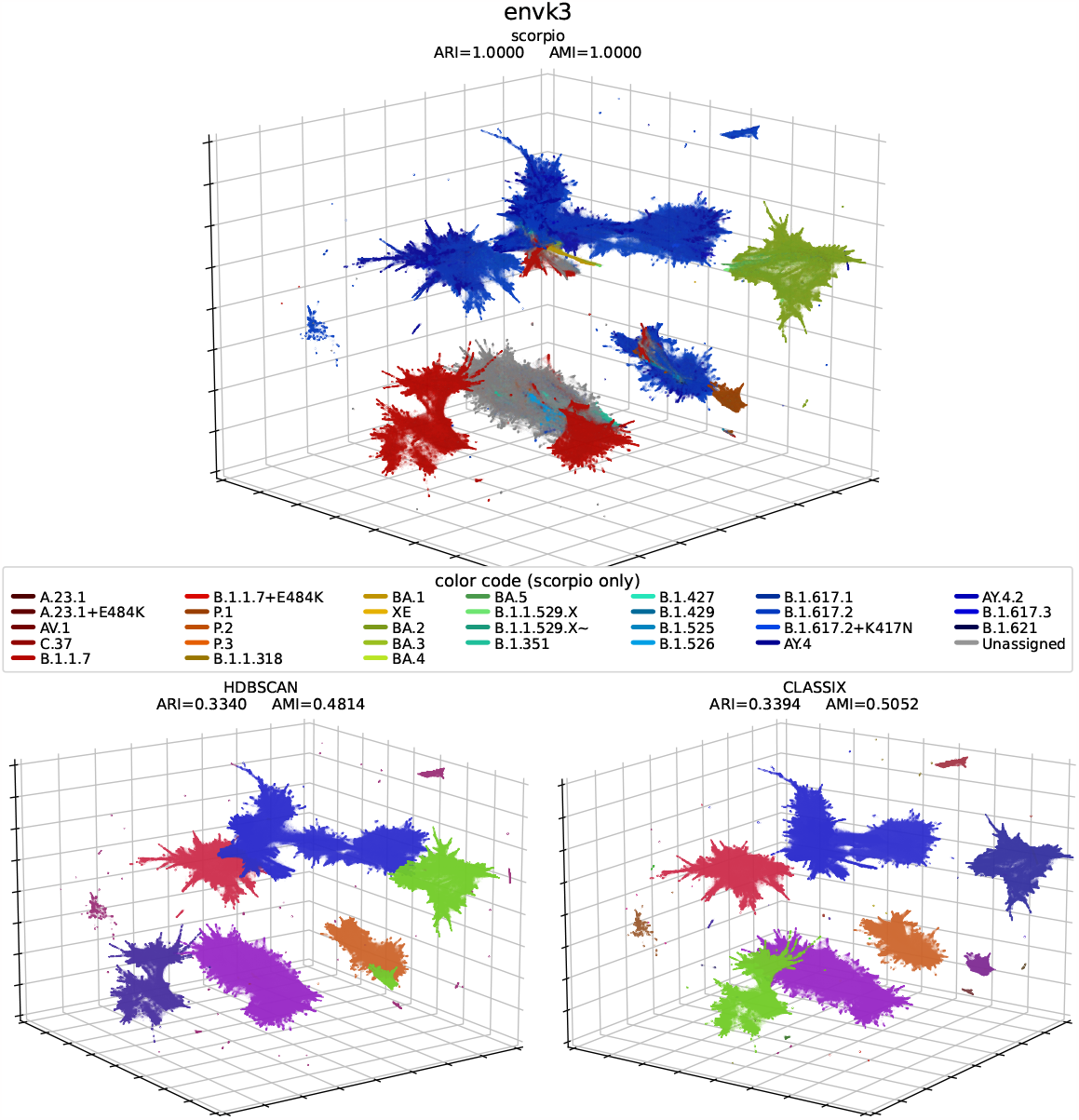
3d PaCMAP projection (parameters: *NB* = 51, *MN* = 0.25 and *FP* = 4) of the whole GISAID dataset extracting the *env* feature [27] (top) coloured by Scorpio labelling [26], (bottom-left) clustering by HDBSCAN [21], (bottom-right) clustering by the novel CLASSIX algorithm [5]. In the bottom figures the colour code is arbitrary, intended just to show the unsupervised differentiation the clustering algorithms perform. The formation of well defined clusters show the potential of these tools to help us gain insight on phylogenetical relations of vast amounts of sequences in an relatively computationally inexpensive fashion. Click on the hyperlinks in this caption to see the interactive 3d plots.

**Figure S2:**
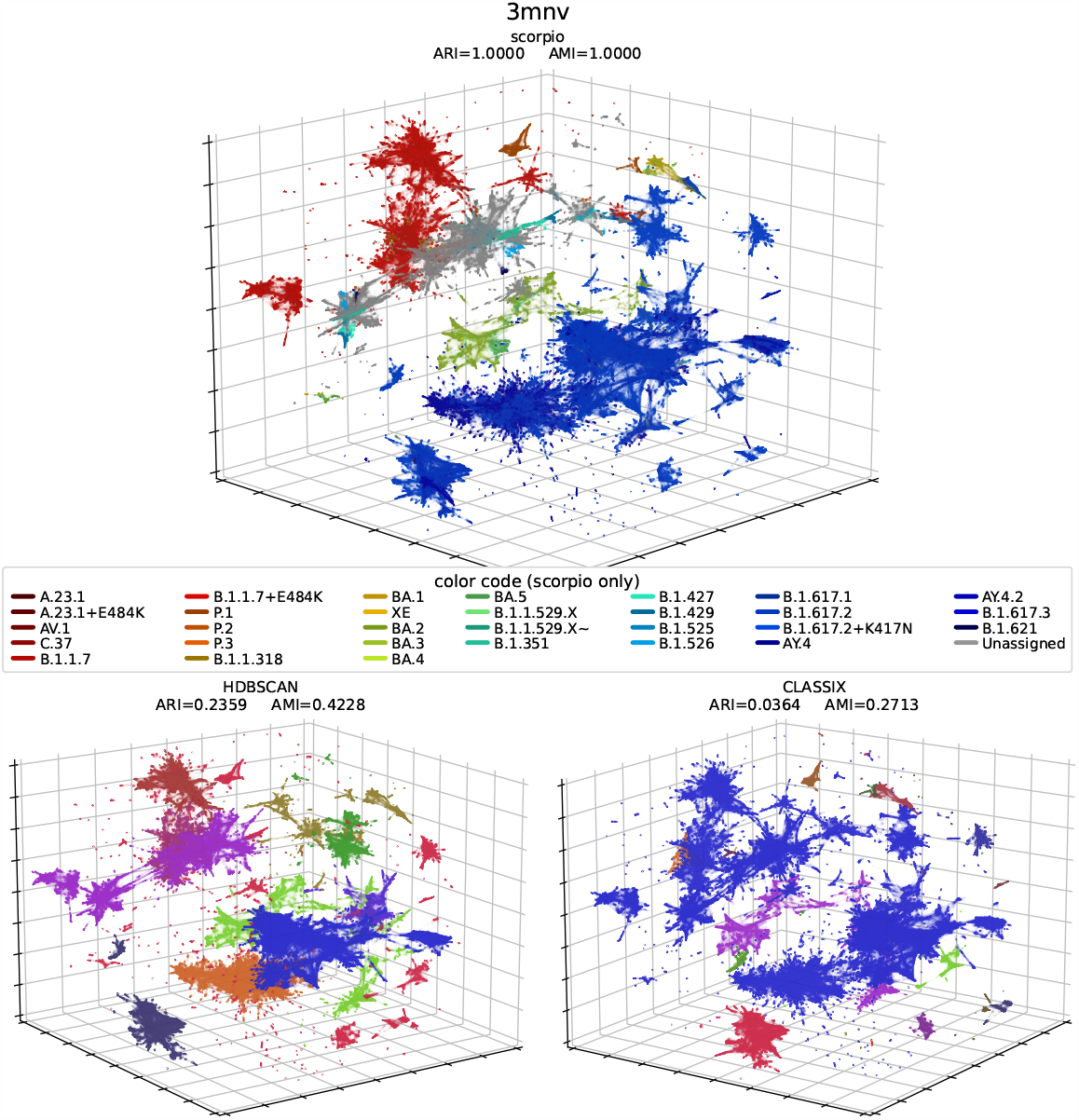
3d PaCMAP (parameters: *NB* = 51, *MN* = 0.25 and *FP* = 2) projection of the whole GISAID dataset extracting the *3mnv* feature [40] (top) coloured by Scorpio labelling [26], (bottomleft) clustering by HDBSCAN [21], (bottom-right) clustering by the novel CLASSIX algorithm [5]. In the bottom figures the colour code is arbitrary, intended just to show the unsupervised differentiation the clustering algorithms perform. The formation of well defined clusters show the potential of these tools to help us gain insight on phylogenetical relations of vast amounts of sequences in an relatively computationally inexpensive fashion. Click on the hyperlinks in this caption to see the interactive 3d plots.

**Figure S3:**
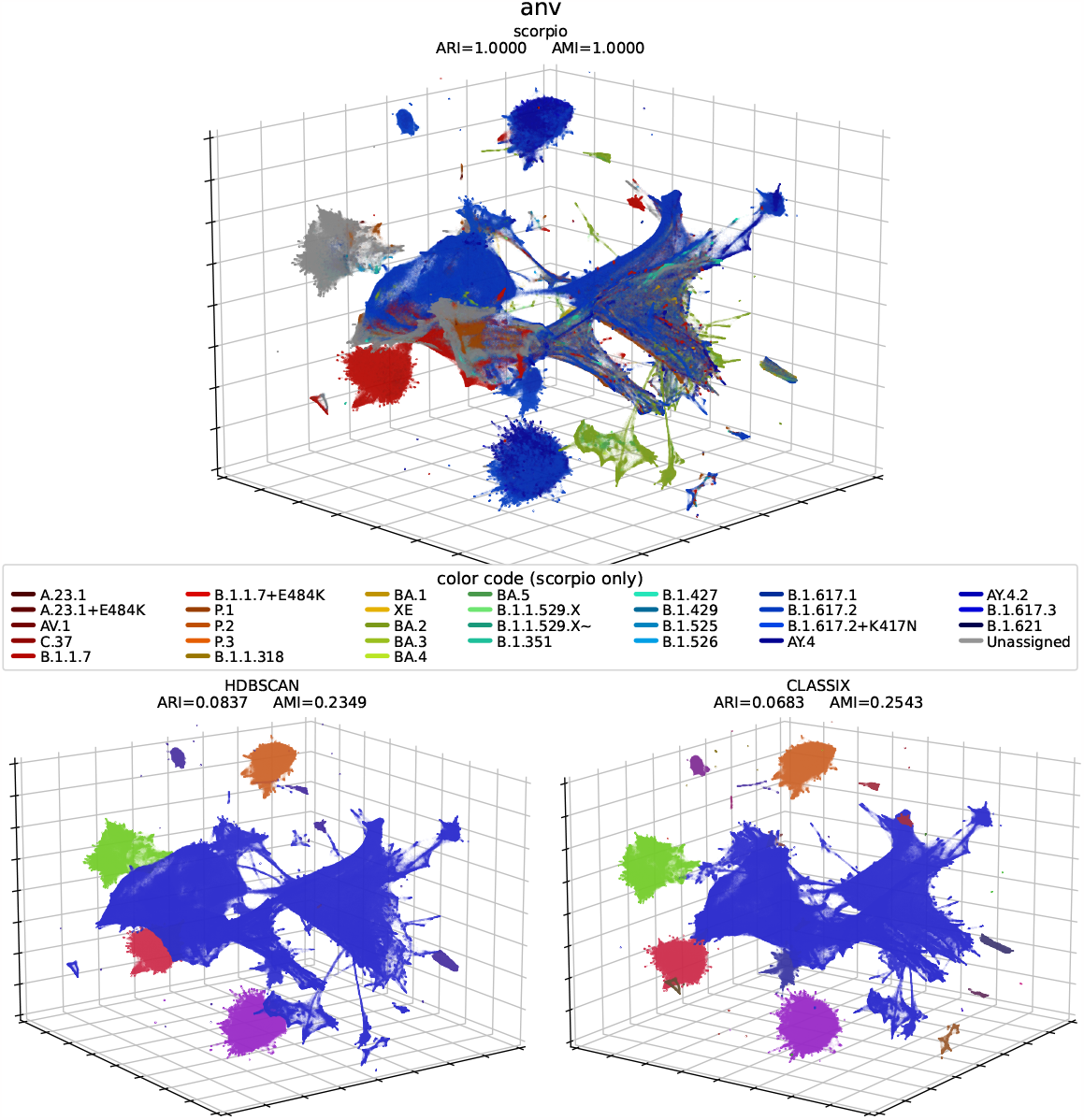
3d PaCMAP projection (parameters: *NB* = 51, *MN* = 1 and *FP* = 4) of the whole GISAID dataset extracting the *anv* feature [8] (top) coloured by Scorpio labelling [26], (bottom-left) clustering by HDBSCAN [21], (bottom-right) clustering by the novel CLASSIX algorithm [5]. In the bottom figures the colour code is arbitrary, intended just to show the unsupervised differentiation the clustering algorithms perform. The formation of well defined clusters show the potential of these tools to help us gain insight on phylogenetical relations of vast amounts of sequences in an relatively computationally inexpensive fashion. Click on the hyperlinks in this caption to see the interactive 3d plots.

**Figure S4:**
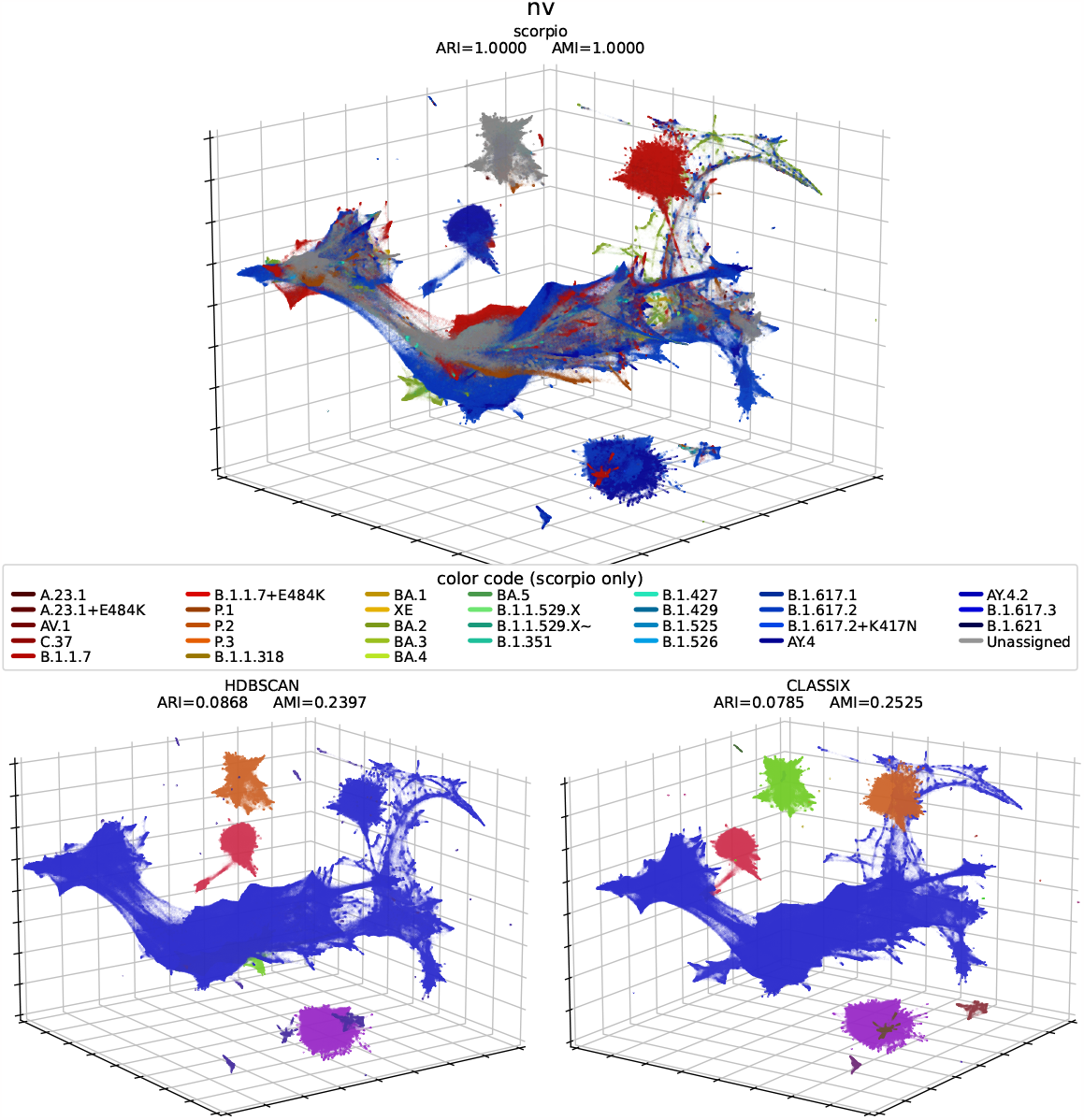
3d PaCMAP projection (parameters: *NB* = 51, *MN* = 1 and *FP* = 2) of the whole GISAID dataset extracting the *nv* feature [7] (top) coloured by Scorpio labelling [26], (bottom-left) clustering by HDBSCAN [21], (bottom-right) clustering by the novel CLASSIX algorithm [5]. In the bottom figures the colour code is arbitrary, intended just to show the unsupervised differentiation the clustering algorithms perform. The formation of well defined clusters show the potential of these tools to help us gain insight on phylogenetical relations of vast amounts of sequences in an relatively computationally inexpensive fashion. Click on the hyperlinks in this caption to see the interactive 3d plots.

**Figure S5:**
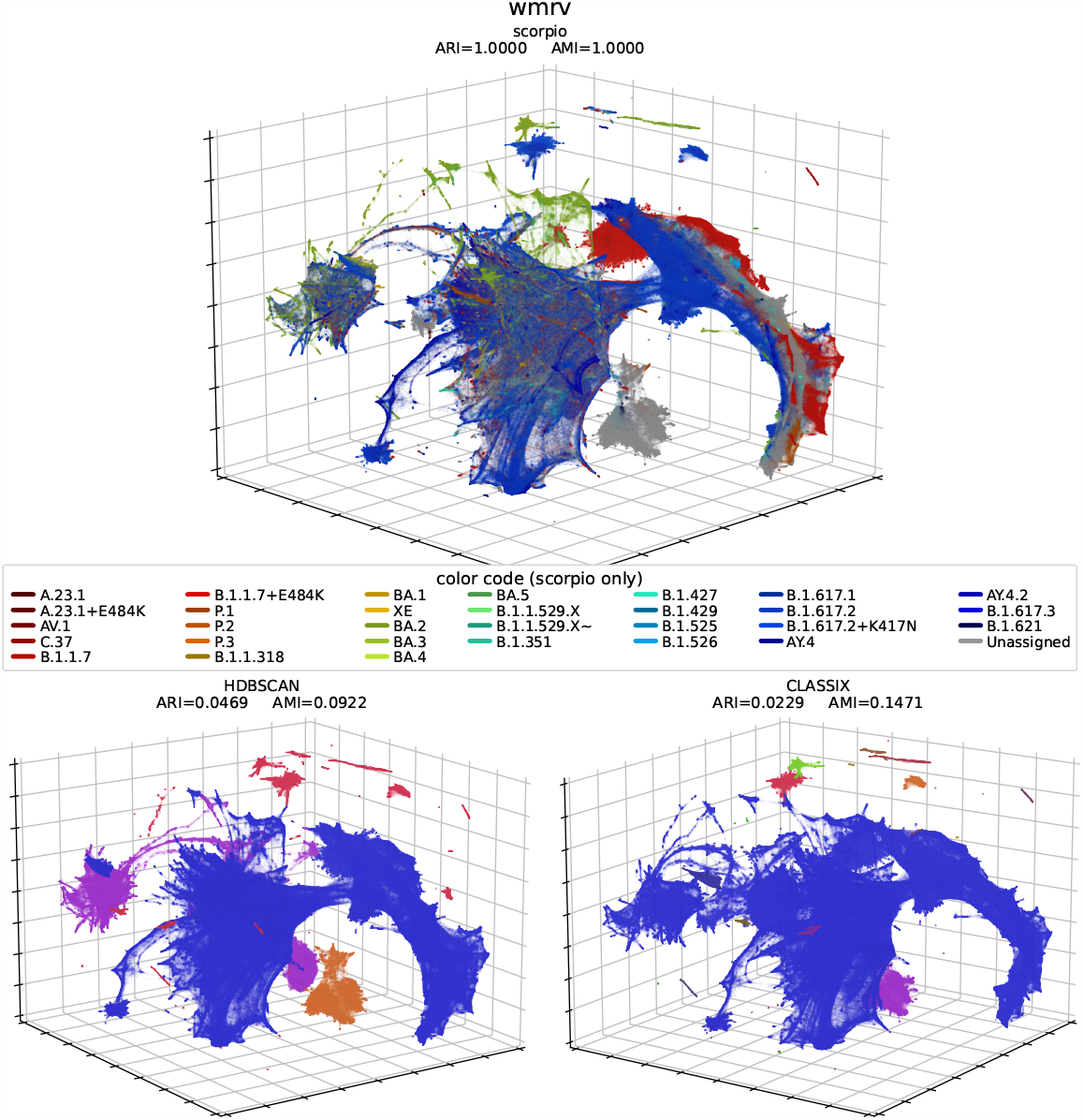
3d PaCMAP projection (parameters: *NB* = 51, *MN* = 50 and *FP* = 4) of the whole GISAID dataset extracting the *wmrv* [19] feature (top) coloured by Scorpio labelling [26], (bottomleft) clustering by HDBSCAN [21], (bottom-right) clustering by the novel CLASSIX algorithm [5]. In the bottom figures the colour code is arbitrary, intended just to show the unsupervised differentiation the clustering algorithms perform. The formation of well defined clusters show the potential of these tools to help us gain insight on phylogenetical relations of vast amounts of sequences in an relatively computationally inexpensive fashion. Click on the hyperlinks in this caption to see the interactive 3d plots.

**Figure S6:**
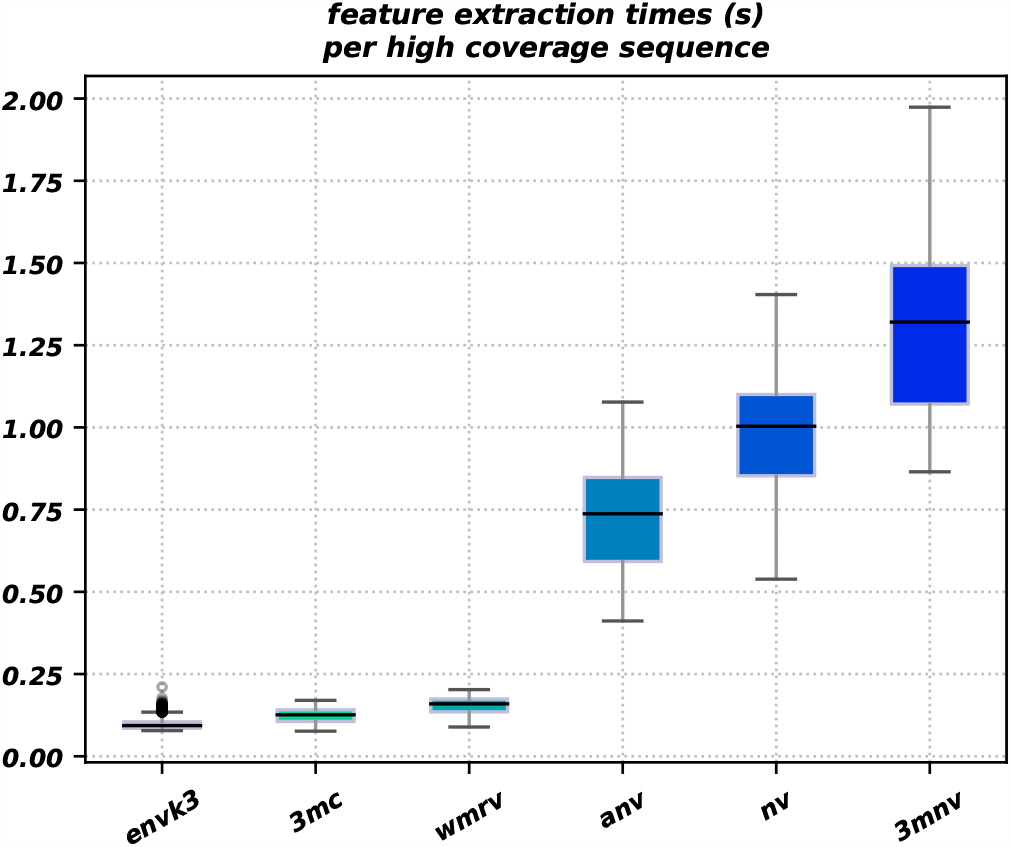
In this plot it is shown an estimation of the processing time for the extraction of each featuring method.

**Table S1:**
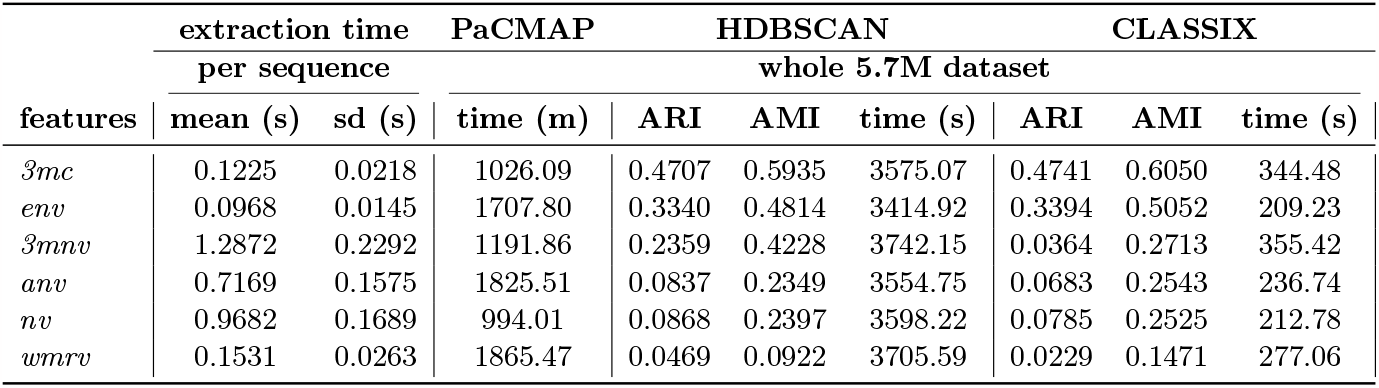
Summary table for all word-statistics features, including: feature extraction process mean time and its standard deviation per high coverage sequence, time to produce the low PaCMAP dimensional projection, and ARI and AMI evaluation for both clustering algorithms.

